# Non-invasive Assessment of Liver Disease in Rats Using Multiparametric Magnetic Resonance Imaging: A Feasibility Study

**DOI:** 10.1101/261032

**Authors:** Anna M. Hoy, Natasha McDonald, Ross J. Lennen, Matteo Milanesi, Amy H. Herlihy, Timothy J. Kendall, William Mungall, Michael Gyngell, Rajarshi Banerjee, Robert L. Janiczek, Philip S. Murphy, Maurits A. Jansen, Jonathan A. Fallowfield

## Abstract

**Background & Aims:** Non-invasive quantitation of chronic liver disease using multiparametric MRI has the potential to refine clinical care pathways, trial design and preclinical drug development. The aim of the study was to evaluate the use of multiparametric liver MRI in experimental rat models of chronic liver disease. ***Methods***: Liver injury was induced in male wild-type Sprague-Dawley rats using 4 or 12 weeks carbon tetrachloride (CCl_4_) intoxication and 4 or 8 weeks methionine and choline deficient (MCD) diet. Liver MRI was performed using a 7.0 Tesla small animal scanner at baseline and specified timepoints after liver injury. Multiparametric liver MRI parameters (T1 mapping, T2* mapping and proton density fat fraction (PDFF)) were correlated with gold standard histopathological measures, assessed by a blinded expert adjudicator. One-way analysis of variance with Tukey’s post-hoc test was used to assess differences between groups and Spearman’s rank-order correlation to examine associations.

**Results:** Mean hepatic T1 increased significantly in rats treated with CCl_4_ for 12 weeks compared to controls (1122±78 ms vs. 959±114 ms; d=162.7, 95% CI (11.92, 313.4), *P*=0.038) and correlated strongly with histological collagen content (r_s_=0.717, *P*=0.037). In MCD diet-treated rats, hepatic PDFF correlated strongly with histological fat content (r_s_=0.819, *P*<0.0001), steatosis grade (r_s_=0.850, *P*<0.0001) and steatohepatitis score (r_s_=0.818, *P*<0.0001). Although there was minimal histological iron, progressive fat accumulation in MCD diet-treated liver significantly shortened T2*.

**Conclusions:** In preclinical models, quantitative MRI markers correlated with histopathological assessments, especially for fatty liver disease. Validation in longitudinal studies is required.

**Key points:**

- Non-invasive quantitative measures of chronic liver injury are needed for stratification and monitoring of disease in clinical practice and in drug development
- We adapted and applied a clinically-relevant non-contrast multiparametric MRI protocol in well-established preclinical liver disease models, for simultaneous quantitation of hepatic fat, fibro-inflammatory injury and iron
- Multiparametric liver MRI was feasible at 7.0 Tesla in experimental rat models and imaging parameters correlated with gold standard histopathological assessments, especially characteristics of fatty liver disease
- Further development of multiparametric liver MRI could refine drug development strategies and impact upon trial design and clinical care pathways

## Introduction

Chronic liver disease (CLD) caused by obesity, excessive alcohol consumption and viral hepatitis represents a significant and increasing healthcare and socio-economic burden worldwide. Non-alcoholic fatty liver disease (NAFLD) is now the commonest form of CLD with a global prevalence of 25% (1). Despite advances in our understanding of the pathogenic mechanisms that underlie NAFLD and liver fibrosis in general, there are still no FDA-approved drug therapies. Important therapeutic targets and promising drug candidates have been identified in preclinical models, but reliable triage of efficacious therapies requires optimization and standardization of drug development strategies and consensus on acceptable endpoints (2). In particular, reproducible and scalable quantitative non-invasive markers of chronic liver injury that permit stratification and longitudinal monitoring of disease could refine preclinical testing, increase efficiency of trial design, and transform clinical care pathways.

Current clinical trials of potential anti-NASH or antifibrotic drugs are anchored to histopathological assessments of disease progression. However, liver biopsy is hampered by invasiveness and sampling variability and is unpopular with patients. Magnetic resonance imaging (MRI) has emerged as a promising new modality for improved understanding and characterization of *in vivo* liver pathophysiology. It is a versatile technique whereby structural and functional MRI can be performed in a single multiparametric scan session, depicting changes associated with inflammation, fibrosis, steatosis, siderosis, oxygenation, and tissue microstructure. Quantitative MRI parameters characterize these processes directly in the liver, rather than using downstream/upstream metabolic analytes. Compared to histopathology, MRI is non-invasive and avoids sampling bias by characterizing the whole organ with high spatial resolution. The MRI proton density fat fraction (PDFF) is a highly accurate and reproducible technique for the assessment of steatosis and can detect changes in hepatic fat as small as 1% (3). Magnetic resonance elastography (MRE) is a phase-contrast MR technique that measures liver stiffness as a surrogate of fibrosis. MRE has high accuracy for the diagnosis of advanced liver fibrosis and cirrhosis, but it is not yet known whether it is sufficiently sensitive or dynamic for the longitudinal monitoring of fibrosis progression/regression (4). Liver*MultiScan*^TM^ (Perspectum Diagnostics, Oxford, UK) uses non-contrast multiparametric MRI to quantify hepatic fibro-inflammatory injury (iron-corrected T1 mapping), steatosis (PDFF) and iron content (T2* mapping). In patients, Liver*MultiScan*^TM^ can accurately quantify liver disease (5-7), may predict clinical outcomes (8), and is now being adopted into clinical trial protocols as a surrogate endpoint. We hypothesized that multiparametric MRI could be developed and applied in experimental models of NASH and liver fibrosis using a small animal MR scanner system, with potential future utility in preclinical drug development studies whilst also addressing the principles of the 3Rs for humane animal research.

## Materials and Methods

### Animal models

Animal experiments were conducted with local and Government ethical approval and in compliance with the Use of Animals in Scientific Procedures Act 1986. Experiments were designed and implemented according to NC3R ARRIVE guidelines. Animals were housed under standard conditions (12:12h light-dark cycle, temperature 23±2°C, with free access to chow and water), 3 rats per cage, and were acclimatized to the room for 1 week before the beginning of the experiments.

### Rat carbon tetrachloride (CCl_4_) model

Chronic carbon tetrachloride (CCl_4_) intoxication in rodents is a reproducible, well-characterized model that induces liver fibrosis (after ~4 weeks) and cirrhosis (after ~12 weeks). Liver injury was induced in 250-300g male wild-type Sprague-Dawley rats by twice-weekly intraperitoneal (i.p.) injection of 0.2 mL/100g CCl_4_ (Sigma-Aldrich, Gillingham, UK) in a 1:1 ratio with sterile olive oil vehicle (OO; Sigma-Aldrich, Gillingham, UK), for either 4 or 12 weeks (n=6 per group). Littermate control rats (n=4 per group) were injected with an identical volume of OO. Animals underwent MRI at the end of the 4 or 12 week treatment course under inhalational general anesthesia (1–2% isoflurane in oxygen) without recovery. Body weight, liver and spleen weights were recorded and liver tissue and blood were collected for histological and biochemical analysis. One of the 12 week CCl_4_-treated animals was euthanized prematurely due to poor health and excluded from analysis.

### Methionine and choline deficient diet model

The methionine and choline deficient (MCD) diet is a frequently used rodent model of NASH. After 4 weeks of MCD diet, aminotransferase levels are elevated and hepatic histology shows florid centrilobular hepatocellular steatosis, whilst 8 weeks of MCD diet results in NASH and early fibrosis. Steatohepatitis was induced in 250-300g male wild-type Sprague-Dawley rats by feeding a lipogenic MCD diet (Research Diets, New Jersey, USA) for 4 or 8 weeks (n=6 per group). Littermate control rats received standard chow (n=4 per group). Animals underwent MRI at the end of the 4 or 8 week treatment course under inhalational general anesthesia (1–2% isoflurane in oxygen) without recovery. Body weight, liver and spleen weights were recorded and liver tissue and blood were collected for histological and biochemical analysis. In a separate experiment, rats were fed an MCD diet for a total of 4 weeks. After 2 weeks, animals were randomized to treatment with either 100 mg/kg/day of cilostazol (a phosphodiesterase type-3 inhibitor previously shown to reduce steatohepatitis in experimental NAFLD models (9)) or vehicle by daily gavage from week 2 to week 4 (n=6 per group). Animals underwent MRI at baseline and after 4 weeks.

### *In vivo* MR imaging

All animals were continually monitored for vital signs and to maintain depth of anesthesia. Respiration rate was monitored by detection of breathing motion by compression of a respiratory sensor placed in contact with the abdomen and core temperature by a rectal thermometer.

### Multiparametric liver MRI

Imaging studies were performed in anesthetized rats using a 7.0 Tesla preclinical MRI scanner (Agilent Technologies, Santa Clara, CA, USA), with a 72 mm internal diameter volume coil. At each timepoint, the following scans were run: two gradient-echo scans with arrayed echo times (repetition time (TR) 75 ms, 4 averages, flip angle 20º), one scan with 4 in-phase echoes (1.01, 2.02, 3.03, 4.04 ms) and one with 4 out-of-phase echoes (1.52, 2.53, 3.54, 4.55 ms), to measure fat using a 3-point Dixon method, and T2* using the 4 in-phase echoes. An adapted gradient echo (fast low angle shot (FLASH)) Look-Locker sequence was used to measure T1 (acquired with 5 inversion times ranging from 270 to 5100 ms), 2 averages, flip angle 8º, echo train length 32). All scans were respiratory gated with a field of view (FOV) 60 mm × 60 mm, 2 mm thick single slice, 128 x 128 matrix. The pulse sequence was based on previous work (10), but was adapted to use an inversion recovery module rather than a saturation module. The axial slice was positioned to cover an abdominal region containing primarily liver. Depending on respiration and anesthesia levels, the full set of scans including shimming and localization were completed in 40-50 min.

### MR image analysis

Data was exported in phase and magnitude format. T2* was determined using the in-phase gradient echo data (11). Pixels in which the magnitude failed to exceed the background noise by 2-fold (11) were excluded from the analysis. A combination of the in-phase and out-of-phase gradient echo data were used to calculate proton density fat and water maps using the extended 3-point Dixon approach described by Glover and Schneider (12). The PDFF was calculated from the fat (F) and water (W) maps as

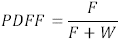

T1 maps were calculated from the Look-Locker data as described previously (13). From each of the T1, T2* and PDFF maps three regions of interest (ROI) over the whole liver were collected per timepoint per animal. Vessels and non-liver regions were not included in ROIs.

To investigate the potential effect of parenchymal fat on hepatic T1, we also performed localized proton magnetic resonance spectroscopy (1H-MRS) using a point resolved spectroscopy (PRESS) sequence at two distinct locations depicting different signal intensity on the T2-weighted structural scan. The voxel dimensions were 4 × 4 × 4 mm, TR = 3 s and echo time (TE) = 23 ms. One signal average was used.

### Serum liver enzyme measurement

Serum alanine aminotransferase (ALT), was measured as described previously (14) utilizing a commercial kit (Alpha Laboratories Ltd., Eastleigh, UK). Aspartate aminotransferase (AST) was determined by a commercial kit (Randox Laboratories, Crumlin, UK). Total bilirubin was determined by the acid diazo method (15) using a commercial kit (Alpha Laboratories Ltd., Eastleigh, UK). All assays were adapted for use on a Cobas Fara centrifugal analyzer (Roche Diagnostics Ltd., Welwyn Garden City, UK). Within-run precision of these assays was calculated and expressed as coefficient of variation (CV) < 4%, while intra-batch precision was CV < 5%.

### Liver histology assessments

Liver tissue was fixed in 4% phosphate-buffered formaldehyde and embedded in paraffin, or in OCT and frozen on dry ice and stored at −80 ° C until histological analysis. Five-micron sections were stained for activated hepatic stellate cells by alpha-smooth muscle actin (α-SMA) (Sigma-Aldrich, Gillingham, UK) and interstitial fibrosis by picrosirus red, as described previously (16). Histology was assessed independently by an expert liver histopathologist, blinded to treatment allocation. Livers were staged for fibrosis by the injury-independent modified Ishak score (scale 0-6; Supplementary Table 1). Both CCl_4_ and MCD diet induce steatosis and lobular necroinflammation, modeling histological features characterizing NAFLD. In human disease, this is assessed by application of the NAFLD activity score (NAS). The same features of steatotic inflammatory injury in the rodent models were scored on a scale from 0 to 8 (steatosis 0-3, lobular inflammation 0-3 and hepatocyte ballooning 0-2) on hematoxylin and eosin (H&E) stained sections using the same criteria (and referred to as the NAFLD (model) Activity Score (N(m)AS) to reflect this application (17). Hepatic iron (ferric form) was detected using Perls’ Prussian Blue and scored using the Scheuer method (18) with grade 0 being negative and grades 1, 2, 3, and 4 representing increasing amounts of stainable iron. Frozen sections were stained with Oil Red O to assess liver fat content. Sections were analyzed using an AxioScan Z1 slide scanner at x20 magnification and ImageJ software (NIH, Bethesda, USA) to quantify the collagen proportionate area (CPA), α-SMA and Oil Red O staining, assessed by a blinded assessor.

### Statistical analysis

Statistical analysis was performed using GraphPad Prism Version 6.0 (GraphPad Software Inc., San Diego, USA). One-way analysis of variance (ANOVA) with Tukey’s post-hoc test was used to assess differences between groups. Spearman’s rank-order correlation (r_s_) was used to examine associations. For all tests, a *P*-value <0.05 was taken to indicate statistical significance. Graphical data is presented as mean±standard deviation (SD).

## Results

### Assessment of hepatic fibro-inflammatory injury in rat CCl_4_ model

To evaluate multiparametric MRI for the assessment of hepatic fibro-inflammatory injury in a preclinical setting we used the rat CCl_4_ model. Compared to OO controls, animals receiving CCl_4_ had higher serum transaminase levels and mild to moderate liver inflammation after both 4 and 12 weeks CCl_4_ (Figure S1). Hepatic fibrogenic activity increased progressively, with 12-fold greater α-SMA expression after 12 weeks in CCl_4_-treated rats compared to controls (*P*=0.006) (Figure S1). Hepatic collagen content (assessed by CPA) was increased, greater than 3-fold, after both 4 weeks (*P*=0.043) and 12 weeks (*P*=0.013) CCl_4_ treatment relative to control (Figure 1A).

**Figure 1.**
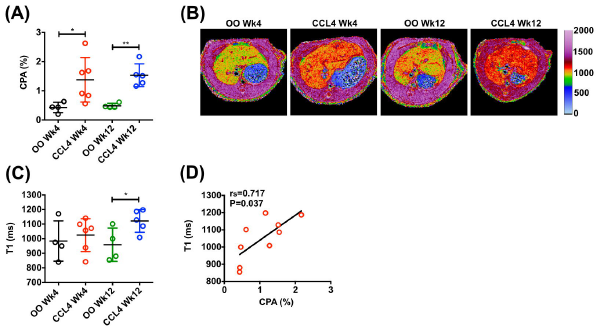
MRI assessment of hepatic fibro-inflammatory injury in rat CCl_4_ model. **A)** Quantification of hepatic collagen content (% area of picrosirius red staining). **B)** Representative examples of transverse liver MR T1 relaxation maps. **C)** T1 relaxation quantification in rats receiving CCl_4_ or olive oil (OO) for 4 and 12 weeks. **D)** Correlation between hepatic collagen content and T1 in rats receiving CCl_4_ and OO for 12 weeks. Data presented as mean±SD, analyzed by unpaired t-test (\**P*<0.05, \*\**P*<0.01). Spearman (r_s_) correlation coefficient was used to examine correlations.

After 4 weeks, liver T1 was 1025±112 ms in CCl 4 rats compared to 984±138 ms in OO controls (d=40.68, 95% CI (−141.7, 223.1), *P*=0.621) (Figure 1B,C). There was a significant difference in mean liver T1 in 12 week CCl_4_ rats (1122±78 ms) compared to controls (959±114 ms; d=162.7, 95% CI (11.92, 313.4), *P*=0.038) (Figure 1B,C). There was a strong positive correlation between liver T1 values and collagen content after 12 weeks CCl_4_ (advanced fibrosis) (r_s_=0.717, *P*=0.037) (Figure 1D) but this correlation was absent after 4 weeks of CCl_4_ (r_s_=-0.079, *P*=0.838) (early bridging fibrosis).

To explore the influence of fat on hepatic T1, we used 1H-MRS to re-analyse two distinct areas identified by different signal intensity on the T2-weighted structural scan. Although one area had a much higher fat content, the T1 values were similar (Figure S2).

### Assessment of hepatic steatosis and steatotic inflammatory injury in preclinical liver injury models

To explore the potential utility of multiparametric MRI as a non-invasive preclinical diagnostic method for the assessment and staging of NAFLD and evaluation of anti-NASH treatments, we used the rat MCD diet model. Rats fed a MCD diet developed florid micro- and macro-vesicular steatosis (Figure S3). The proportionate area of Oil Red O-stained fat in the liver was <0.5% in control rats, but in comparison was grossly elevated after 4 weeks MCD diet (22.9±10.9%; *P*=0.004) and 8 weeks MCD diet (30.2±13.2%; *P*=0.002) (Figure 2B) with no significant difference between both timepoints.

**Figure 2.**
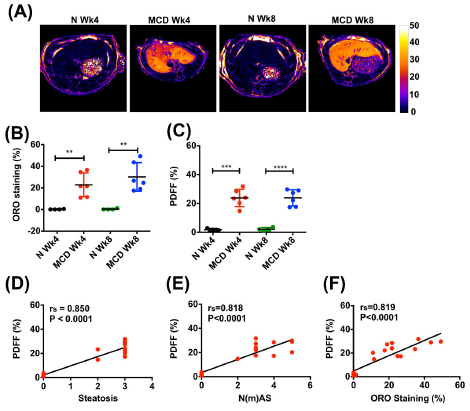
MRI assessment of hepatic steatosis and steatotic inflammatory injury in rat MCD diet model. **A)** Representative examples of transverse liver MR PDFF maps. **B)** Quantification of Oil red O (ORO) staining of hepatic fat. **C)** Quantification of hepatic proton density fat fraction (PDFF). Correlations between hepatic PDFF and **D)** histologically assessed steatosis, **E)** steatotic inflammatory injury (N(m)AS) score and **F)** ORO staining in rats receiving MCD or control diet (N) for 4 and 8 weeks. Data presented as mean±SD, analyzed by unpaired t-test (\*\**P*<0.01, \*\*\**P*<0.001, \*\*\*\**P*<0.0001). Spearman (r_s_) correlation coefficient was used to examine correlations.

Quantification of hepatic fat by MRI (mean of 3 ROIs) showed a highly significant increase in PDFF after both 4 and 8 weeks of MCD diet, reaching 23.81±6.00% and 23.84±5.64% compared with control animals where PDFF was 1.73±0.94% (*P*=0.0005) and 2.09±1.16% ( *P*<0.0001), respectively (Figure 2A,C). The hepatic PDFF strongly correlated with fat proportionate area (r_s_=0.819, *P*<0.0001) (Figure 2F). Furthermore, hepatic PDFF correlated with both histological stage of steatosis and steatotic inflammatory injury (r_s_=0.850, *P*<0.0001 and r_s_=0.818, *P*<0.0001 respectively) (Figure 2D,E).

In contrast to animals receiving MCD diet, CCl_4_ treated rats displayed a smaller increase in histological liver fat compared with controls after 4 weeks (8.94±6.48% vs. 0.26±0.19%, *P*=0.03) and 12 weeks (9.11±9.6% vs 0.30±0.13%, *P*=0.11) (Figure 3B). Quantification of hepatic fat by MRI (Figure 3A) showed an increase in PDFF compared to controls after 4 weeks CCl_4_ (3.86±1.83% vs 0.91±0.93%, *P*=0.016) and 12 weeks CCl_4_ (3.58±0.33% vs. 1.85±0.87%, *P*=0.0044) (Figure 3C). Moreover, PDFF was strongly correlated with histological grading of steatosis (Oil Red O staining) and steatotic inflammatory injury (r_s_=0.74, *P*=0.003 and r_s_=0.84, *P*<0.0001 respectively) (Figure 3D,E).

**Figure 3.**
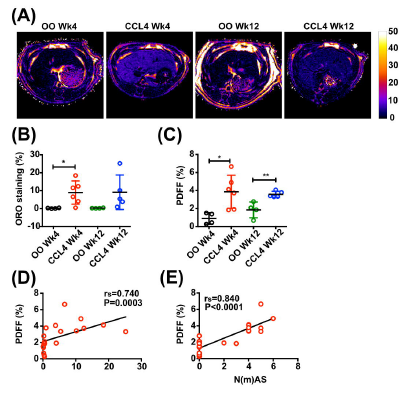
MRI assessment of hepatic steatosis and steatohepatitis in rat CCl_4_ model. **A)** Representative examples of transverse liver MR PDFF maps. **B)** Quantification of Oil red O (ORO) staining of hepatic fat. **C)** Quantification of hepatic proton density fat fraction (PDFF). Correlations between hepatic PDFF and **D)** histologically assessed ORO staining and **E)** steatotic inflammatory injury (N(mAS) score in rats receiving CCl_4_ or olive oil (OO) for 4 and 12 weeks. Data presented as mean±SD, analyzed by unpaired t-test (* *P*<0.05, \*\**P*<0.01). Spearman (r_s_) correlation coefficient was used to examine correlations.

It had previously been shown that administration of cilostazol for 16 weeks reduced hepatic steatosis, inflammation and fibrosis in rats fed a choline-deficient, l-amino acid-defined (CDAA) diet or a high-fat high-calorie diet. We tested our MRI methodology in rats that were fed MCD diet for 4 weeks plus treatment with cilostazol or vehicle control between weeks 2-4. Animals underwent multiparametric MRI at baseline and after 4 weeks. On histological examination, cilostazol treatment for 2 weeks only reduced hepatic steatosis by approximately 5% (d=-5.19%, 95% CI (−13.91, 3.53), *P*=0.21), with a non-significant reduction in N(m)AS score (Figure 4 A,B). Nevertheless, in a drug intervention setting multiparametric MRI methods performed similarly (Figure 4C,D) and, importantly, strong correlations were observed between the histological assessment of steatotic inflammatory injury and MRI measures (PDFF; r_s_=0.675, *P*=0.019) (Figure 4E).

**Figure 4.**
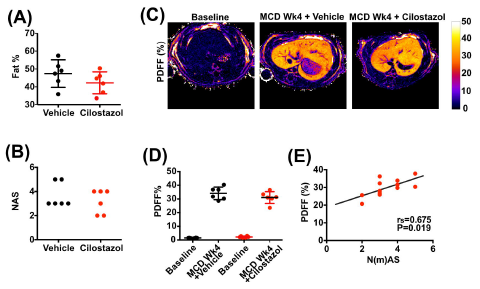
MRI assessment of hepatic steatosis and steatohepatitis in cilostazol treated rats on MCD diet. Sprague-Dawley rats received MCD diet for 4 weeks plus cliostazol or vehicle for the last 2 weeks of the model. Animals were scanned at baseline and after 4 weeks. **A)** Quantification of hepatic fat by morphometry based on H&E staining. **B)** Steatotic inflammatory injury (N(m)AS) score. **C)** Representative examples of transverse liver MR proton density fat fraction (PDFF). **D)** Quantification of hepatic PDFF. **E)** Correlation between PDFF and steatotic inflammatory injury (N(m)AS) score. Data presented as mean±SD, analyzed by unpaired t-test. Spearman (r_s_) correlation coefficient was used to examine correlations.

### Assessment of hepatic iron content in preclinical liver injury models

Hepatic iron may have a pathogenic role in NAFLD and, intriguingly, NAFLD itself may affect iron metabolism (19). From a technical standpoint, both hepatic steatosis and fibrosis can have confounding effects on T2* relaxation times. We showed histologically that CCl_4_ and MCD diet induced injury models were not associated with significant accumulation of iron in the liver (Figure S1, Figure S3), but we did observe variable effects on T2* in these models (Figure 5A). Consistent with the lack of tissue iron staining, there was no change in liver R2* (1/T2*) in CCl_4_ treated rats compared with controls (*P*>0.05) (Figure 5B). However, we observed a significant increase in R2* (1/ T2*) in MCD diet-treated rat livers compared with controls at 4 weeks (177.9±46.7s ^−1^ vs. 76.2±7.9s ^−1^, *P*<0.0001) and 8 weeks (213.6±22.5s ^−1^ vs. 83.6±12.1s ^−1^, *P*<0.0001) (Figure 5C). Moreover, there was a strong positive association between R2* and PDFF (r_s_=0.790, *P*<0.001) (Figure 5D), indicating that steatosis had a cofounding effect on T2* relaxation time in this model.

**Figure 5.**
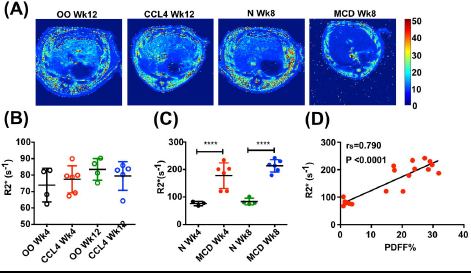
MRI assessment of T2*/R2* relaxation in preclinical liver injury models. A) Representative examples of transverse liver MR T2* maps. **B)** Quantification of R2* (1/T2*) in rats receiving CCl_4_ or olive oil (OO) for 4 and 12 weeks. **C)** Quantification of R2* (1/T2*) in rats receiving MCD or control diet (N) for 4 and 8 weeks. **D)** Correlation between R2* and PDFF in animals receiving CCl_4_ or OO injections and MCD or control diets. Data presented as mean±SD, analyzed by unpaired t-test (\*\*\*\**P*<0.0001). Spearman (r_s_) correlation coefficient was used to examine correlations.

## Discussion

Despite the identification of a large number of therapeutic targets from preclinical studies, there are currently no licensed drug treatments for liver fibrosis or for NASH. Drug development has been hindered by a reliance on liver biopsy and a lack of robust validated non-invasive biomarkers to diagnose and monitor changes in liver disease over time or in response to treatment. Multiparametric MRI using Liver*Multiscan*^TM^ has shown promise as a clinical diagnostic tool and surrogate endpoint in therapeutic trials, especially in the assessment of NAFLD. Our objective was to develop and apply this non-invasive, contrast-free multiparametric MRI methodology in preclinical liver injury models that are commonly used in mechanistic and drug evaluation studies.

Here we showed that our multiparametric liver MRI protocol was feasible at 7.0 Tesla in experimental rats and scanning was completed within a realistic time frame (~40 minutes, compared to ~85 minutes for a combined preclinical MRE and collagen molecular imaging protocol) (20). Both hepatic T1 and PDFF changed with increasing duration of liver injury and were strongly correlated with histological assessments of fibrosis and steatosis/steatotic inflammatory injury, respectively. Unfortunately, there was only a minimal effect of a short treatment course of cilostazol in the MCD diet model and even histological endpoints were not significantly altered within this abbreviated timeframe. Although a number of MRI techniques have been evaluated individually in animal models to assess hepatic fibrosis (19), iron content (20) or fat (21), multiparametric MRI captures all of these parameters in a single contrast-free scan protocol. In a previous study in mice with fatty liver, non-invasive MR assessments (1H-MRS, PDFF and modified Dixon) had a higher correlation with liver triglyceride content than digital image analysis of Oil Red O stained liver sections (21). The authors concluded that MRI was the preferred method for fat quantification. The performance of Liver*Multiscan*^TM^ in our experimental rat models was broadly similar to that observed in patients with CLD (r_s_=0.68, *P*<0.0001 for fibrosis; r_s_=0.89, *P*<0.001 for steatosis; r_s_=-0.69, *P*<0.0001 for siderosis) (5).

Our findings indicate that multiparametric liver MRI could potentially refine the way we currently approach testing of therapeutic candidates in preclinical models. Firstly, multiparametric MRI could be used as a quantitative and objective method to stratify liver disease severity before exposing animals to a drug treatment. It is known that animals maintained on an experimental lipogenic diet develop NAFLD at differing rates and of varying severity and, similarly to human disease, fibrosis accumulates heterogeneously within the liver. A pretreatment liver (wedge) biopsy protocol was previously used to categorize NAFLD mice into three groups (steatosis without fibrosis, NASH with fibrosis, cirrhosis), in order to address the inherent and potentially confounding issue of variability in the study population (22). Imaging with MRI could be used in a similar manner, especially where accurate stratification could permit a more precise evaluation of therapies that modulate different components of NAFLD pathogenesis (i.e. anti-steatotic, anti-inflammatory, antifibrotic), but would have the advantages of being non-invasive and sampling the whole liver. Secondly, multiparametric MRI could potentially be used longitudinally to evaluate the effect of an antifibrotic or anti-NASH intervention in individual experimental animals, allowing each to serve as its own control, and thereby allowing determination of the distribution of effects in an experimental population (22). This approach would also address the 3Rs (refinement, reduction, replacement) of humane animal experimentation.

We acknowledge the limitations of this study. Firstly, this was an initial small feasibility study to determine whether a multiparametric MRI protocol could be successfully scaled down and applied in rats with chronic liver injury within a realistic timeframe. Our findings require further validation and confirmation of reproducibility, in order to use this test in longitudinal/interventional studies. Secondly, technical refinements to the protocol may increase robustness and reliability. The small organ size of rodents necessitates the use of high-field MR imaging systems to obtain adequate temporal and spatial resolution and poses major challenges to imaging due to shortened relaxation times and accentuated field inhomogeneity. Additionally, hepatic T1 values could potentially be influenced by the fat fraction, thus confounding its use as a quantitative marker of hepatic fibro-inflammatory injury. To explore this, we re-analyzed one rat using 1H-MRS that clearly had two distinct areas in the liver, one with a much higher fat content. However, the T1 value for both areas was the same. Nevertheless, in contrast to histological collagen quantification, we found that T1 mapping was not sufficiently sensitive to discriminate between normal liver and mild fibrosis (after 4 weeks CCl_4_). This remains the Achilles’ heel of non-invasive serum and imaging biomarkers in the clinical setting. Interestingly, a recent study in rats with liver fibrosis showed that MRE was only sensitive for the detection of late-stage fibrosis, whereas a collagen-specific molecular imaging probe could only distinguish between earlier stages of fibrosis (20). The differing sensitivity of various MRI methods indicates complementary staging capabilities and the potential for developing clinically-relevant multiparametric composite MRI models. The utility of T2* as a quantifiable marker for iron overload is well established (23) and T2* is currently the clinical standard for the non-invasive assessment of iron overload. However, low tissue iron content in our rat liver injury models meant that we were unable to confirm the expected inverse relationship between increasing iron load and decreased T2 and T2* relaxation time in liver tissue. Furthermore, as demonstrated in this study, T2* is influenced by a number of factors independent of tissue iron concentration. In particular, we observed a significant shortening of T2* in the lipogenic MCD diet model, despite very low hepatic iron levels. This finding is consistent with previous human studies that confirm a short apparent T2* of fat (24, 25).

Using multiparametric MRI to quantify liver injury in experimental animal models could be a powerful new tool to assist in the development of new therapies for CLD, and is directly translatable to clinical practice.

## Acknowledgments

The work was carried out on a preclinical MRI scanner within the Edinburgh Imaging Facility, University of Edinburgh.

## Abbreviations

ALT/AST: Alanine/aspartate aminotransferase
ANOVA: One-way analysis of variance
CCl_4_: Carbon tetrachloride
CDAA: Choline-deficient, l-amino acid-defined
CLD: Chronic liver disease
CPA: Collagen proportionate area
CV: Coefficient of variation
H&E: Hematoxylin and eosin
i.p.: Intraperitoneal
MCD: Methionine choline deficient
MRE: Magnetic resonance elastography
MRI: Magnetic resonance imaging
MRS: Magnetic resonance spectroscopy
ms: Millisecond
N(m)AS: NAFLD (model) Activity Score
NAFLD: Non-alcoholic fatty liver disease (NAFLD)
NASH: Non-alcoholic steatohepatitis (NASH)
OO: Olive oil
ORO: Oil Red O
PDFF: Proton density fat fraction
PRESS: Point RESolved Spectroscopy
ROI: Region of interest
SD: Standard deviation
α-SMA: Alpha-smooth muscle actin

